# Regulation of the SOS response and homologous recombination by an integrative and conjugative element

**DOI:** 10.1101/2024.10.11.617942

**Authors:** Alam García Heredia, Alan D. Grossman

## Abstract

Integrative and conjugative elements (ICEs) are mobile genetic elements that transfer between bacteria and influence host physiology and promote evolution. ICE*Bs1* of *Bacillus subtilis* modulates the host DNA damage response by reducing RecA filament formation. We found that the two ICE*Bs1*-encoded proteins, RamT and RamA that modulate the SOS response in donors also function in recipient cells to inhibit both the SOS response and homologous recombination following transfer of the element. Expression of RamT and RamA caused a decrease in binding of the host single strand binding protein SsbA to ssDNA. We found that RamA interacted with PcrA, the host DNA translocase that functions to remove RecA from DNA, likely functioning to modulate the SOS response and recombination by stimulating PcrA activity. These findings reveal how ICE*Bs1* can modulate key host processes, including the SOS response and homologous recombination, highlighting the complex interplay between mobile genetic elements and their bacterial hosts in adaptation and evolution.

## Introduction

The genetic landscape and evolution of bacteria can be shaped by horizontal gene transfer. This process is often mediated by mobile genetic elements (MGEs), including conjugative plasmids and integrative and conjugative elements (ICEs) that can transfer from one bacterial cell to another. In addition to genes that are needed for their own transfer, these conjugative elements often carry genes that provide phenotypes to host cells [reviewed in (Johnson & Grossman, 2015; Delavat et al., 2017)].

ICEs typically reside integrated in the host chromosome. Some ICEs target specific sites for integration, and others are capable of integrating at multiple locations throughout the chromosome [reviewed in (Johnson & Grossman, 2015; Delavat et al., 2017)]. Unlike plasmids, ICEs encode their own recombinases and do not rely on homologous recombination for integration. When activated in a donor, or upon transfer to a new host, many ICEs undergo autonomous rolling circle replication. The timing of ICE integration is regulated to ensure that autonomous rolling circle replication ceases either prior to or concurrently with integration (McKeithen-Mead & Grossman, 2023). In contrast, plasmids are generally maintained extrachromosomally, although some can integrate into the host chromosome, typically by homologous recombination. For example, the F plasmid of *Escherichia coli* can integrate into the *E. coli* chromosome by homologous recombination between insertion sequences present in both the plasmid and the host chromosome [reviewed in (Vos et al., 2024)].

Conjugative elements typically transfer as single-stranded DNA (ssDNA) during bacterial conjugation (Johnson & Grossman, 2015; Delavat et al., 2017). ssDNA is also generated during rolling circle replication of an ICE. This ssDNA can be recognized by host cells as a signal of DNA damage, serving as a substrate for RecA protein. RecA binds to ssDNA, forming nucleoprotein filaments that subsequently activate the SOS response, a global stress response to DNA damage [reviewed in (Baharoglu & Mazel, 2014; Maslowska et al., 2019)]. Some conjugative plasmids have evolved mechanisms to inhibit RecA activity and suppress the SOS response (Petrova et al., 2009; Al Mamun et al., 2021).

ICE*Bs1* from *Bacillus subtilis* contains two genes (*ramT* and *ramA*) that function to modulate the host SOS response when the element is active (McKeithen-Mead et al., 2024). Previous genetic analyses revealed that both proteins inhibit the SOS response, and RamT can also activate it in the absence of RamA (McKeithen-Mead et al., 2024). The model for these interactions is RamT activates the ability of RamA to inhibit SOS, and RamT independently of RamA, stimulates SOS.

Here, we explored how of RamT and RamA function to reduce the SOS response. We found that, in addition to their known role in decreasing SOS in donor cells (McKeithen-Mead et al., 2024), both RamT and RamA reduce the SOS response in recipient cells after transfer of the element DNA during conjugation. Additionally, we found that RamT and RamA inhibit homologous recombination. It is known that RecA filament formation on ssDNA is inhibited by single-stranded binding proteins by direct competition (Shan et al., 1997; Manfredi et al., 2008). We found that RamT functions to decrease the association of the single-stranded DNA binding protein SsbA with ssDNA, possibly explaining its role as an activator of SOS when present alone. We also found that RamA interacts directly with the DNA translocase PcrA, likely enhancing its known ability to dismantle pre-existing RecA filaments (Anand et al., 2007; Park et al., 2010; Carrasco et al., 2022). These mechanisms enable ICE*Bs1* to modulate the host’s DNA damage response and homologous recombination when ICE*Bs1* DNA is single-stranded.

## Results

### RamT and RamA decrease levels of the SOS response in recipient cells following conjugative DNA transfer

When ICE*Bs1* is induced in a population of host cells (donors), it elicits the SOS response in a subpopulation of those cells. The absence of *ramT* and *ramA* increase this response, demonstrating that the normal function of RamT and RamA is to dampen the SOS response (McKeithen-Mead et al., 2024). We sought to determine if *ramT* and *ramA* also affect recipients following transfer of the element to a new host.

#### Assay to detect the SOS response in transconjugants

We used Cre-mediated recombination, with *cre* under control of the SOS-inducible promoter PyneA (PyneA-*cre*), as a readout of SOS in transconjugants. ICE*Bs1* contained PyneA-*cre* and recipients (without ICE*Bs1*) contained a spectinomycin resistance gene (*spc*) with its promoter (P*spc*) flanked by two *loxP* sites and facing opposite *spc* (Fig 1A). During conjugation, if recipients receive ICE*Bs1* (PyneA*-cre*) and undergo SOS, Cre will be produced and catalyze the inversion of P*spc*, which can be detected by qPCR (see Methods). We normalized inversion of P*spc* by total recipients in conjugation experiments to get the proportions of recipient cells that underwent SOS. We measured levels of SOS during conjugation by qPCR instead of spectinomycin resistance since we found that virtually all of the transconjugants eventually became resistant to spectinomycin. This is indicative of some SOS response during growth of the transconjugants, likely due to replication fork arrest that is known to occur in most replication cycles [reviewed in (Lenhart et al., 2012)].

**Figure 1.**
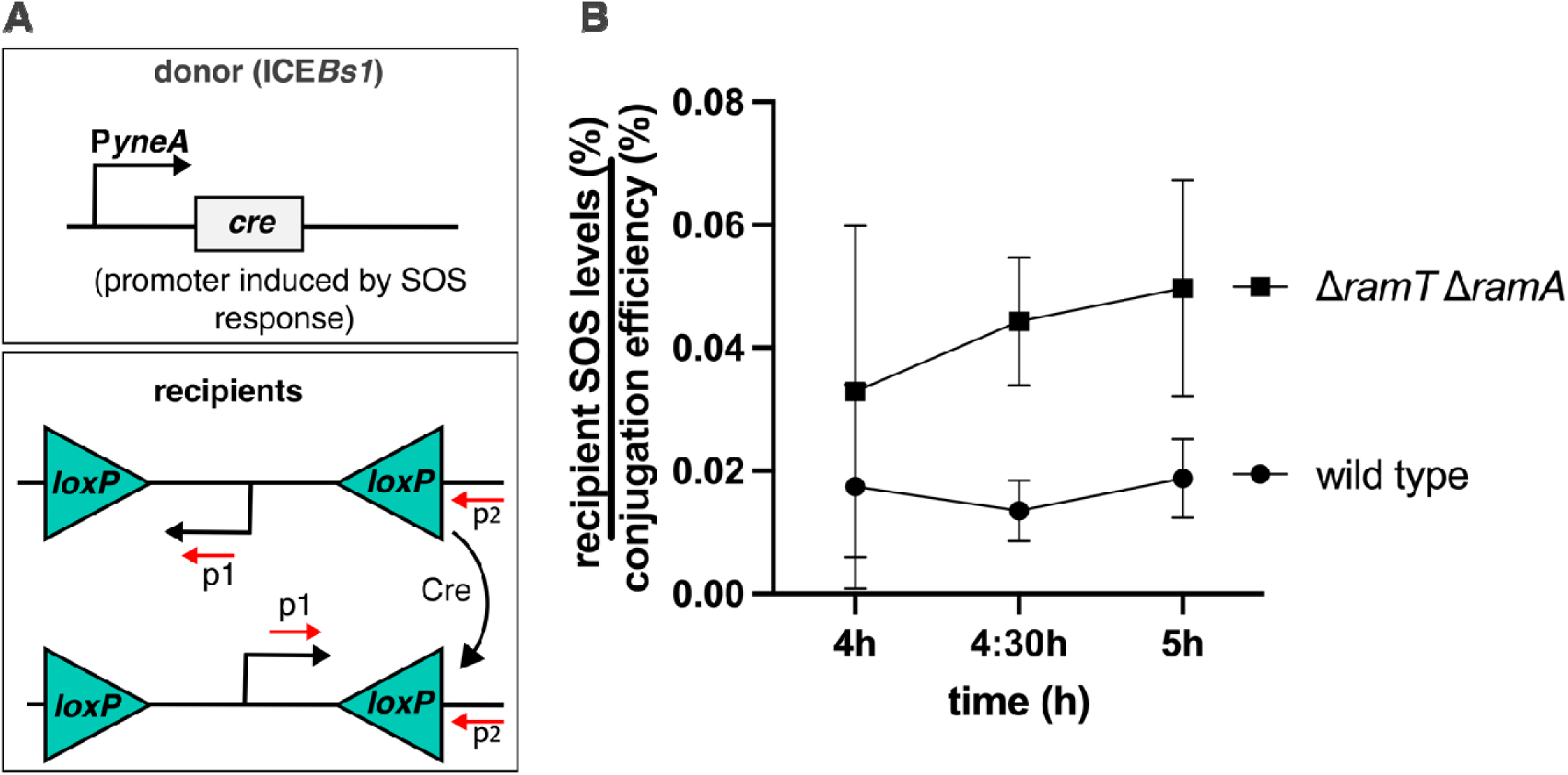
RamT and RamA decrease SOS levels in recipients during conjugation. **A**, experimental setup. ICE*Bs1* carries the Cre recombinase under the SOS promoter P*yneA*. Recipient cells contain a reverse-oriented P*spc* promoter. Primers to P*spc* (p1 and p2) can only amplify the promoter in the correct orientation, which only takes place when a Cre-mediated inversion has occurred. During conjugation, ICE*Bs1* entry to recipients will express the Cre recombinase if recipient cells experience an SOS response. **B**, SOS levels in recipient cells during conjugation. The SOS levels were calculated from individual mattings that were incubated for the indicated time (see Methods). SOS levels shown correspond to the ratio of P*spc* inversion per recipients to the conjugation efficiency per experiment. Wild type ICE*Bs1* triggers an SOS response in recipient cells, which increases ∼ 2-fold when *ramT* and *ramA* are absent.

We note that expression from PyneA (PyneA-*lacZ*; and PyneA-*mNeonGreen*) has been used previously to monitor activation and modulation of SOS by ICE*Bs1* (McKeithen-Mead & Grossman, 2023) (McKeithen-Mead et al., 2024). Also, ICE*Bs1* containing a constitutively expressed *cre* has been used to detect conjugative transfer by qPCR, irrespective of the ability of transconjugants to form colonies (McKeithen-Mead & Grossman, 2023).

#### SOS in a subpopulation of transconjugants

We found that approximately two percent of the recipient cells that received ICE*Bs1* underwent an SOS response during or shortly after conjugation (Fig 1B). Deletion of *ramT* and *ramA* caused an approximately twofold increase in this response after 4 hours of mating compared to wild type ICE*Bs1* (Fig 1B). This amount of SOS in transconjugants is about half of that previously observed in donor cells (McKeithen-Mead et al., 2024). We suspect this apparent difference is due to the transient presence of single-stranded ICE DNA in transconjugants compared to the persistent presence in donors under the conditions previously monitored. Based on these results, we conclude that there is an SOS response in a subset of transconjugants and that RamT and RamA normally function to reduce this response.

### RamT and RamA inhibit homologous recombination

Formation of RecA filaments is an early step in both the SOS response and homologous recombination. Because RamT and RamA inhibit formation of RecA filaments (McKeithen-Mead et al., 2024), we suspected that they might also inhibit homologous recombination. To test this, we measured the conjugation efficiency of an ICE*Bs1* mutant (Δ*int*::*tet*) that is missing the site-specific recombinase and forms transconjugants via homologous recombination (Lee et al., 2007). Recipient cells (AGH533) contained Δ*comK*::*tet*, with *tet* (∼1 kb) providing the region of homology for recombination with ICE*Bs1* Δ*int*::*tet*. The conjugation efficiency of ICE*Bs1* Δ*int*::*tet* (AGH540) was ∼0.001% viable transconjugants per donor (Fig 2B). Deletion of *ramT* and *ramA* (AGH564), increased the conjugation efficiency ∼6-fold (Fig 2B). These results indicate that RamT and RamA decrease homologous recombination. Under the conditions used, the conjugation efficiency of wild type ICE*Bs1* is typically ≥1% transconjugants per donor, about 2-3 orders of magnitude greater than that observed for ICE*Bs1* Δ*int*::*tet*.

**Figure 2.**
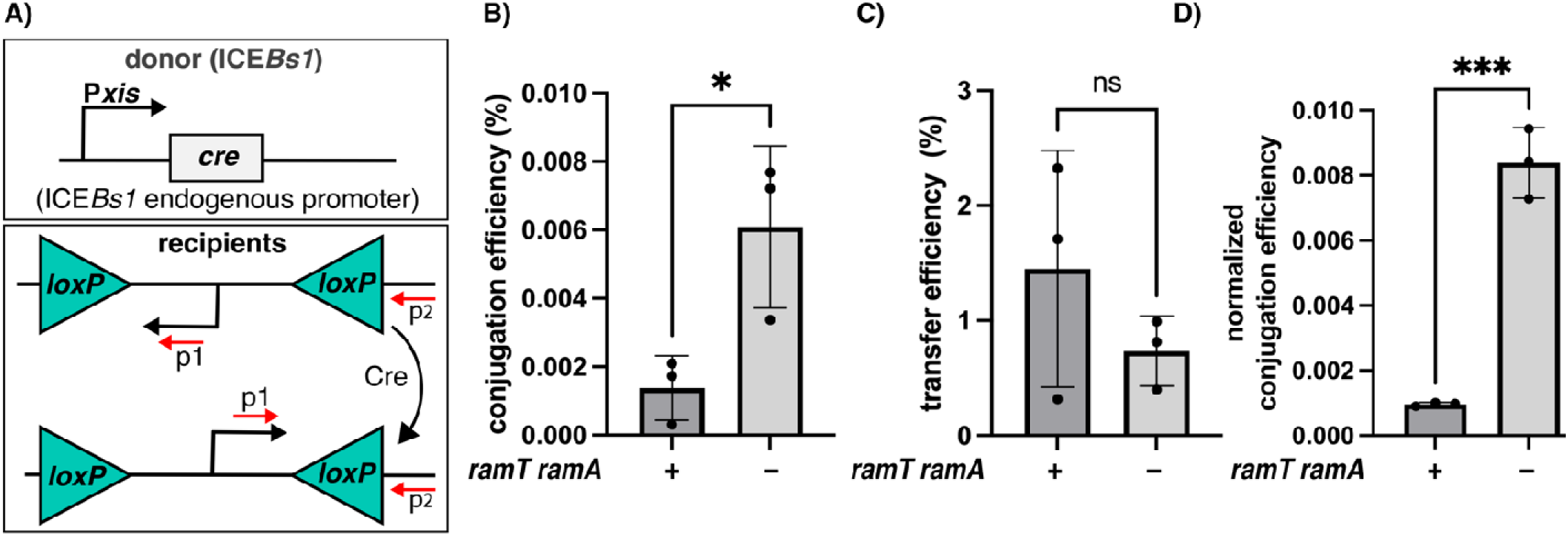
RamT and RamA decrease homologous recombination. **A**, experimental setup. The Cre recombinase is expressed from P*xis* following activation of ICE*Bs1*. Recipient cells contain *loxP* sites flanking P*spc*. Primers to P*spc* (p1 and p2) can only amplify the promoter following Cre-mediated inversion. **B**, mating efficiencies mediated by homologous recombination. Donor cells carrying ICE*Bs1* (Δ*int*::*tet*) either with (AGH540) or without (AGH564) *ramT* and *ramA* were mated with recipient cells (AGH533) containing ∼1 kb homology (*tet*) with the element. Conjugation efficiency was calculated as the ratio of stable transconjugants to initial donor cells (see Methods). **C**, the same strains from panel B were used to determine the transfer efficiency of ICE*Bs1* to recipient cells. qPCR was used to measure Cre-mediated recombination as an indicator for the proportion of recipient cells that received the element during conjugation per total recipients, regardless of integration by homologous recombination or subsequent viability. **D**, the normalized conjugation efficiency was calculated by dividing conjugation efficiency (**B**) by the respective transfer efficiency (**C**) for each experiment. Statistical analysis was performed using Student’s t-test. *p<0.05; ***p<0.0005.

Since generation of transconjugants involves the stable integration of ICE*Bs1* into the chromosome of recipient cells and growth of the resulting integrant, we wanted to consider the number of recipient cells that received the element during conjugation but failed to form stable transconjugants, either due to lack of integration or death of the integrants. Knowing the number of cells that initially receive ICE*Bs1* and the number of stable transconjugants enables us to calculate the number of transconjugants generated per recipient that received ICE*Bs1*. This would also allow for detection of and correction for any variations in conjugation between different cultures.

To calculate the number of recipient cells that received ICE*Bs1*, we used the *cre*-*lox* reporter system, essentially as described previously (McKeithen-Mead & Grossman, 2023). In these experiments, transcription of *cre* was driven by the major ICE*Bs1* promoter P*xis* (Fig 2A) (Methods). Recipients contained the spectinomycin (*spc*) resistance cassette with its promoter in reverse orientation between two *loxP* sites, as described above. Transcription of *cre* would occur in transconjugants after ICE*Bs1* DNA becomes double stranded. Inversion of the *spc* promoter would be indicative of recipient cells that acquired the element, irrespective of successful integration (Fig 2A; Methods). We normalized P*spc* inversion by the total recipient population to extract the numbers of recipients that received ICE*Bs1* during our experiments. We refer to the efficiency of initial acquisition of ICE*Bs1* as the “transfer efficiency”, and this is irrespective of integration or viability. We found that the transfer efficiency of ICE*Bs1*, as determined by qPCR to measure inversion of P*spc*, was a bit more than 1% per donor for wild type (*ramT*^+^ *ramA*^+^; AGH540) and a bit less than 1% per donor for the mutant (Δ*ramT* Δ*ramA*; AGH564; Fig 2C). These differences were not statistically significant (Fig 2C).

We normalized the mating efficiency of ICE*Bs1* containing or lacking *ramT* and *ramA* (Fig 2B) to their respective experimental transfer efficiencies (Fig 2C) within each experiment. We found that the normalized conjugation efficiency of ICE*Bs1* lacking *ramT* and *ramA* was ∼9-fold greater than that of ICE*Bs1* that contained wild type *ramT* and *ramA* (Fig 2D). Together, these results in combination with the effects of RamT and RamA on formation of RecA filaments indicate that the effects of RamT and RamA on the recovery of stable transconjugants (in the absence of the site-specific recombinase Int) is most likely due to effects on homologous recombination, and that RamT and RamA normally function to inhibit homologous recombination.

### RamT decreases binding of the host single strand binding protein SsbA to ssDNA

The essential single-strand DNA binding protein SsbA binds and stabilizes ssDNA during growth and has a complex role in regulating the formation of RecA filaments (Lindner et al., 2004; Meyer & Laine, 1990; Lenhart et al., 2012). SsbA, on its own, inhibits the ability of RecA to filament on ssDNA due to competition (Shan et al., 1997; Manfredi et al., 2008). However, SsbA recruits RecO, which then nucleates RecA into ssDNA to form RecA filaments (Shan et al., 1997; Manfredi et al., 2008). Therefore, SsbA is needed for the SOS response and homologous recombination as it stabilizes ssDNA while also facilitating RecA filament formation through RecO.

Since SsbA can both inhibit and stimulate formation of RecA filaments, we postulated that RamT might modulate SsbA. To test this hypothesis, we used chromatin immunoprecipitation (ChIP) to assess binding of SsbA-GFP to ssDNA (see Methods). We used cells that contained a plasmid without a single-strand origin (*sso*). The absence of the *sso* results in an increase in plasmid ssDNA.

To validate our experimental setup, we first confirmed the preferential association of SsbA-GFP to ssDNA over dsDNA. We used cells containing plasmids that either lack or have an *sso* (*sso^−^* and *sso^+^*, respectively) and immunoprecipitated SsbA-GFP after crosslinking. As expected, we observed approximately 40-fold more immunoprecipitated DNA from the *sso*^-^ plasmid (in strain AGH603) compared to the *sso*^+^ plasmid (in strain AGH604; Fig 3A).

**Figure 3.**
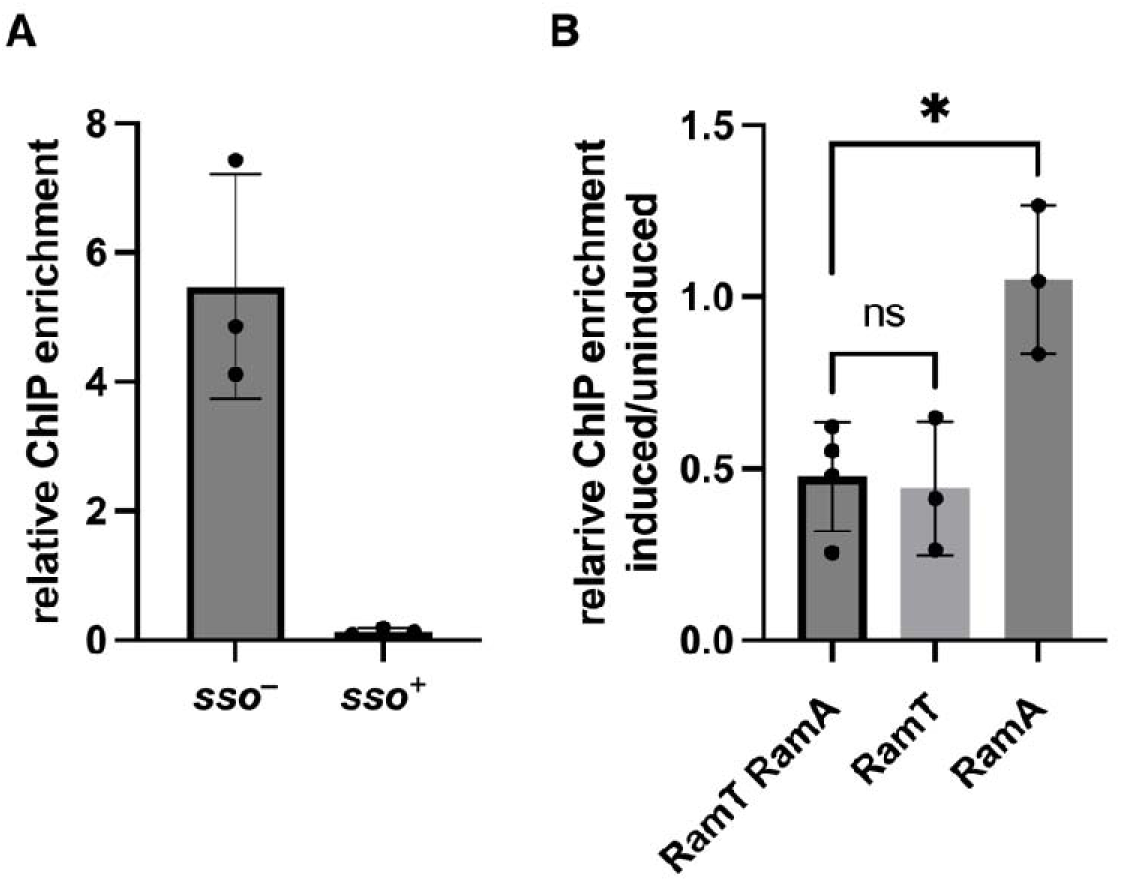
RamT reduces binding of SsbA-GFP to ssDNA. **A)** Following crosslinking, SsbA-GFP was immunoprecipitated in cells containing plasmids (pHP13) with and without an *sso*. SsbA-GFP binds to *sso^−^*∼40-fold more than *sso^+^*, indicating the binding preference towards ssDNA. **B)** RamT and/or RamA were induced in cells expressing SsbA-GFP. SsbA-GFP was then precipitated, and the associated plasmid DNA was quantified using qPCR (see Methods). RamT alone (AGH699) or co-expressed with RamA (AGH603) decreased binding of SsbA-GFP to ssDNA by ∼50%. Expression of RamA alone had no effect on SsbA-GFP binding to ssDNA. Statistical analysis, ANOVA with Tuekeýs test as posthoc. *p < 0.05 indicating statistical significance. Y-axis is the relative amount of plasmid DNA in the ChIP.

We found that RamT but not RamA inhibited binding of SsbA-GFP to ssDNA. In cells that lack ICE*Bs1*, co-expression of *ramT* and *ramA* caused a modest decrease (∼50%) in binding of SsbA-GFP to ssDNA (Fig 3B). Expression of *ramT* alone caused a similar decrease in binding of SsbA-GFP to ssDNA. On the other hand, expression of *ramA* alone had little or no effect on binding of SsbA-GFP to ssDNA. Based on these results, we conclude that RamT reduces binding of SsbA-GFP to ssDNA, either directly or indirectly, and that RamA does not participate in this effect.

The ability of RamT to decrease binding of SsbA-GFP to ssDNA indicates that these two proteins could interact. We used a yeast two-hybrid assay to investigate potential interactions between RamT and SsbA (see Methods). Our results revealed a positive interaction between these two proteins, providing evidence for their ability to associate in the absence of other *B. subtilis* cellular components. However, it’s important to note that this interaction was observed outside the natural cellular context of *B. subtilis* and it remains to be determined whether this interaction occurs under physiological conditions in live *B. subtilis*.

In aggregate, our findings indicate that RamT reduces SsbA binding to ssDNA, and that these proteins likely interact. Theoretically, this reduction of SsbA binding to ssDNA by RamT should facilitate RecA filament formation, explaining the role of RamT as an SOS response activator (McKeithen-Mead et al., 2024). However, both RamT and RamA decrease the amount of SOS levels upon ICE*Bs1* induction (McKeithen-Mead et al., 2024), and when they are co-expressed, SsbA binding is reduced by similar amounts compared to when RamT is expressed alone (Fig 3). This indicates that decreasing binding of SsbA to ssDNA may enable RamA to inhibit SOS induction, consistent with the role of RamT as an SOS inhibitor in the presence of RamA (McKeithen-Mead et al., 2024).

### RamA interacts with PcrA

We started searching for proteins involved in formation of RecA filaments that might interact with RamA. Experiments described below show that RamA interacts with the host DNA translocase PcrA *in vivo*. PcrA dismantles RecA filaments and is essential for host survival (Anand et al., 2007; Park et al., 2010). Based on *in vitro* analyses, SsbA inhibits translocation of PcrA (Carrasco et al., 2022).

We found that RamA interacts with the host DNA translocase PcrA. We constructed a strain (AGH671) carrying functional fusions of RamA (RamA-GFP) and PcrA (PcrA-myc). We immunoprecipitated PcrA-myc with anti-myc antibodies and found that RamA-GFP co-precipitated (Fig 4). RamA-GFP did not precipitate with untagged PcrA (Fig 4), indicating that the co-immunoprecipitation was specific.

**Figure 4.**
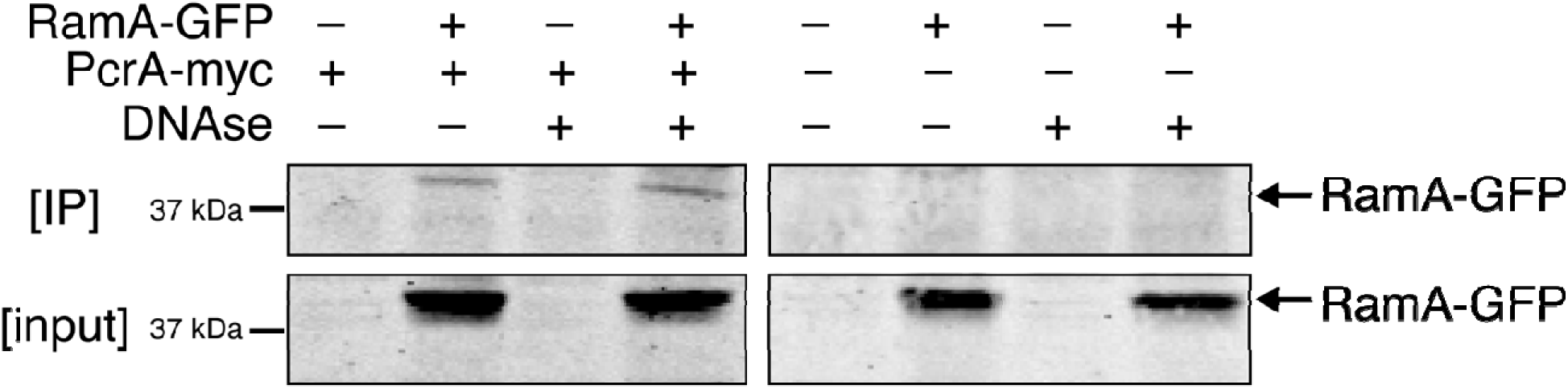
RamA interacts with PcrA. Cells expressing functional fusions to RamA (RamA-GFP) and PcrA (PcrA-myc) were prepared for immunoprecipitation and transferred to filters for Western blotting (see Methods). Anti-myc antibodies were used to immunoprecipitated PcrA-myc. Blots were probed with anti-GFP antibodies to detect RamA-GFP (∼40 kDa) Benzonase (DNase) was added to degrade DNA as indicated. Top, immunoprecipitation. Bottom, input. Arrows indicate RamA-GFP.

We were concerned that the apparent interaction might be due to both proteins being bound to ssDNA and not necessarily near each other or interacting directly, even though crosslinking was not used to detect interactions. If the two proteins were distal to each other on ssDNA, then the apparent interaction would likely be disrupted by DNase treatment. We found that the interaction between PcrA and RamA persisted even after treatment with the DNase benzonase (Fig 4). Based on this result, we infer that PcrA and RamA likely interact directly, or are part of a relatively stable complex. We suspect that this interaction occurs on ssDNA and stimulates PcrA to remove RecA filaments, thereby inhibiting both the SOS response and homologous recombination.

## Discussion

Our work shows that *ramT* and *ramA* of ICE*Bs1* function in transconjugants to inhibit both the SOS response and homologous recombination. Integration of ICE*Bs1* by homologous recombination most likely results in a severe drop in viable transconjugants, despite normal transfer of the element to recipient cells. This drop in viability is most likely due to ongoing autonomous rolling circle replication of ICE*Bs1* after integration and is analogous to the drop in viability when ICE*Bs1* integration by site-specific (Int-mediated) recombination occurs prematurely (McKeithen-Mead & Grossman, 2023). We postulate that by reducing the formation of RecA filaments, ICE*Bs1* not only promotes fitness of its host by reducing growth arrest due to the SOS response, but also promotes faithful integration at its attachment site (*attB*) and reduces possible homologous recombination.

Our findings support a model where the ICE*Bs1*-encoded proteins RamT and RamA work together to reduce the levels of RecA filaments most likely by enhancing the activity of the host DNA translocase PcrA (Fig 5). This mechanism operates through a multi-step process where RamT, either directly or indirectly, decreases binding of SsbA to ssDNA. When bound to ssDNA, SsbA inhibits both RecA filament formation and PcrA DNA translocation. RamT thereby activates both RecA and PcrA by making more ssDNA available.

**Figure 5.**
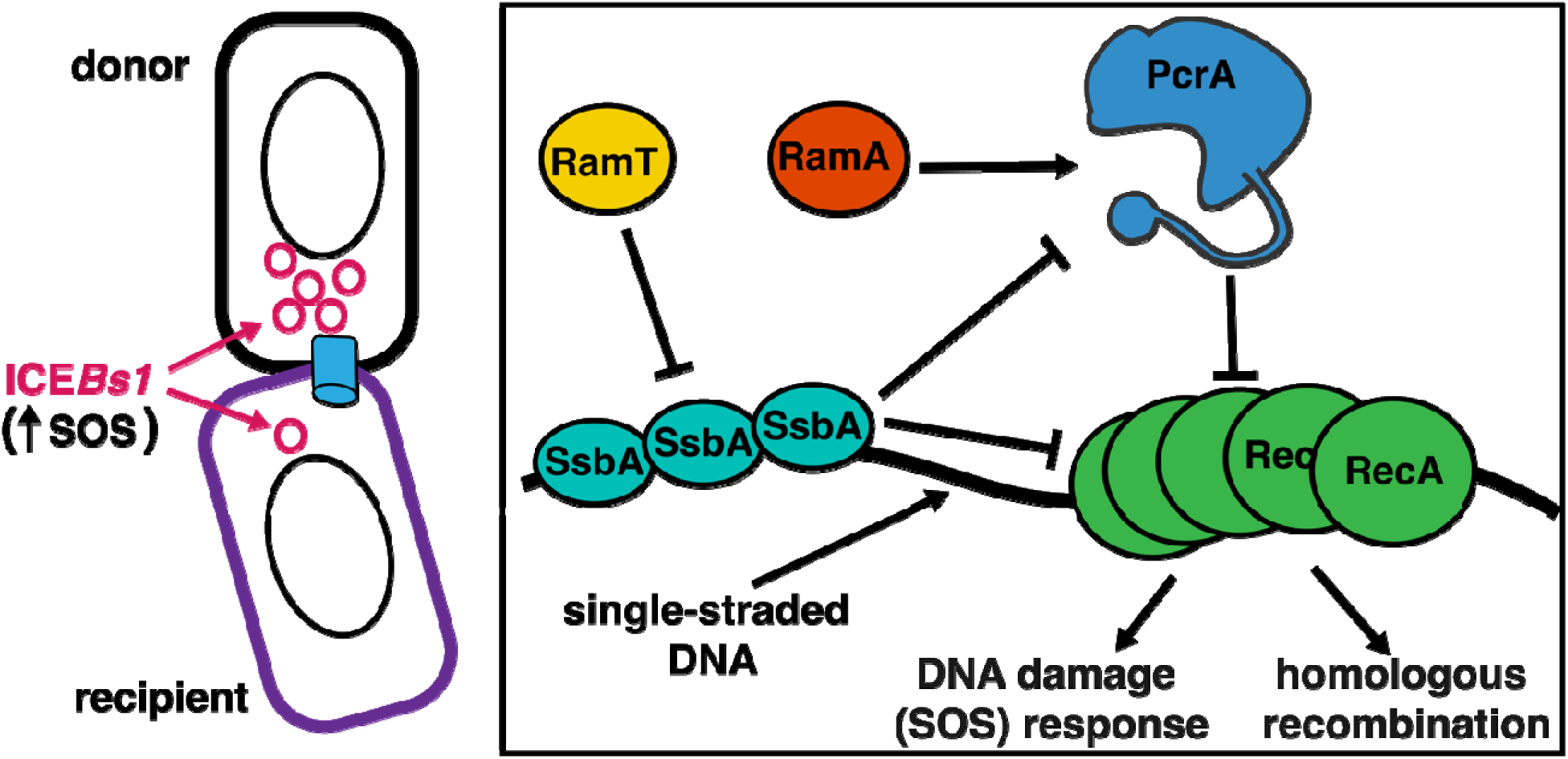
Model of RamT and RamA activity. RamT decreases binding of SsbA to ssDNA. RamA interacts with PcrA and most likely promotes its DNA translocation activity to dismantle RecA filaments.

RamA, known to function only as an inhibitor of the SOS response (McKeithen-Mead et al., 2024), interacts with PcrA and likely enhances its translocation activity to dismantle RecA filaments. Mutant analyses shows that the phenotypes caused by *ramA* depend on the presence of *ramT* (McKeithen-Mead et al., 2024). We believe that the simplest interpretation is that the decreased binding of SsbA caused by RamT enables RamA to stimulate DNA translocation by PcrA. Conversely, in the absence of RamA, the RamT-mediated reduction of SsbA bound to ssDNA likely facilitates the formation of RecA filaments without the enhancement of PcrA activity by RamA, explaining the SOS response activator role of RamT in the absence of RamA. Overall, the regulation of these cellular processes and ultimately the inhibition of RecA filaments reduces the SOS response and likely favors the faithful integration of ICE*Bs1* by site-specific recombination, thereby reducing detrimental effects on the host cell.

The modulation of RecA filament formation and/or the SOS response is a property shared by various mobile genetic elements, including conjugative plasmids and phages, although the mechanisms employed differ significantly. For example, PsiB, encoded by the F plasmid of *E. coli* and other conjugative plasmids bind and inhibit RecA directly (Bagdasarian et al., 1986; Bailone et al., 1988; Petrova et al., 2009; Al Mamun et al., 2021; Baharoglu et al., 2010). In contrast, some bacteriophages take an indirect approach to inhibit the SOS response by targeting proteins other than RecA. For example, bacteriophage λ of *E. coli* encodes proteins that specifically target the RecBCD exonuclease that can affect loading RecA onto DNA (Dillingham & Kowalczykowski, 2008). This is most likely a mechanism to protect phage DNA (Murphy, 1991), but also results in inhibition of formation of RecA filaments. Using a completely different mechanism, the bacteriophage GIL01 of *B. thuringiensis* encodes a protein that inhibits the SOS response by binding to and stabilizing the SOS-repressor LexA (Fornelos et al., 2015).

Our findings highlight how an integrative and conjugative element likely affects the DNA translocase PcrA to reduce RecA filaments. It is interesting to note that ICEs and plasmids that undergo rolling circle replication often use PcrA as a helicase to unwind DNA for replication. We suspect that most ICEs that undergo robust autonomous rolling circle replication likely have mechanisms to inhibit homologous recombination. Further, we postulate that inhibition of RecA by PsiB from conjugative plasmids might also inhibit homologous recombination thereby promoting their dissemination by reducing integration into the chromosome and conversion of cells to Hfrs, which will transfer chromosomal DNA but not the entire integrated element.

## Methods

### Growth media and conditions

*B. subtilis* was grown in LB medium or defined S7_50_ minimal medium with 1% L-arabinose as a carbon source. Serial dilutions were typically done in PBS. For any given experiment, single colonies from the indicated strains were streaked out from frozen stocks (−80°C) into plates with appropriate antibiotics. A single colony was later used to start cultures to be grown to mid-exponential phase. An aliquot was diluted into fresh medium until it reached the culture density appropriate for the experiment. Where indicated, expression of ICE*Bs1*was induced by addition of D-xylose (1% v/v) to overexpress *rapI* from a xylose-inducible promoter (P*xyl*-*rapI*). ICE*Bs1* was also induced using P*spank*-*rapI* and addition of IPTG (2 µM). The antibiotics used include kanamycin (5 μg/ml), tetracycline (8 μg/ml), spectinomycin (100 μg/ml), chloramphenicol (5 μg/ml), and 0.5 μg/ml erthyromycin plus 12.5 μg/ml lincomycin to select for macrolide-lincosamide-streptogramin B (MLS) resistance conferred by *mls* (*ermB*).

### Strains and alleles used

All strains used (Table 1) were derived from JH642 (AG174; (Smith et al., 2014)) and are auxotrophic of phenylalanine and threonine (not referenced on the table). Most strains were constructed by natural transformation and Gibson assembly. In some other cases, ICE*Bs1* was transferred into strains by conjugation. Strains that lack ICE*Bs1* are shown as ICE*Bs1^0^*. Alleles were normally constructed in wild type strains and verified by phenotype, PCR, and sequencing. Alleles were transferred to other strains by transformation.

**Table 1.**
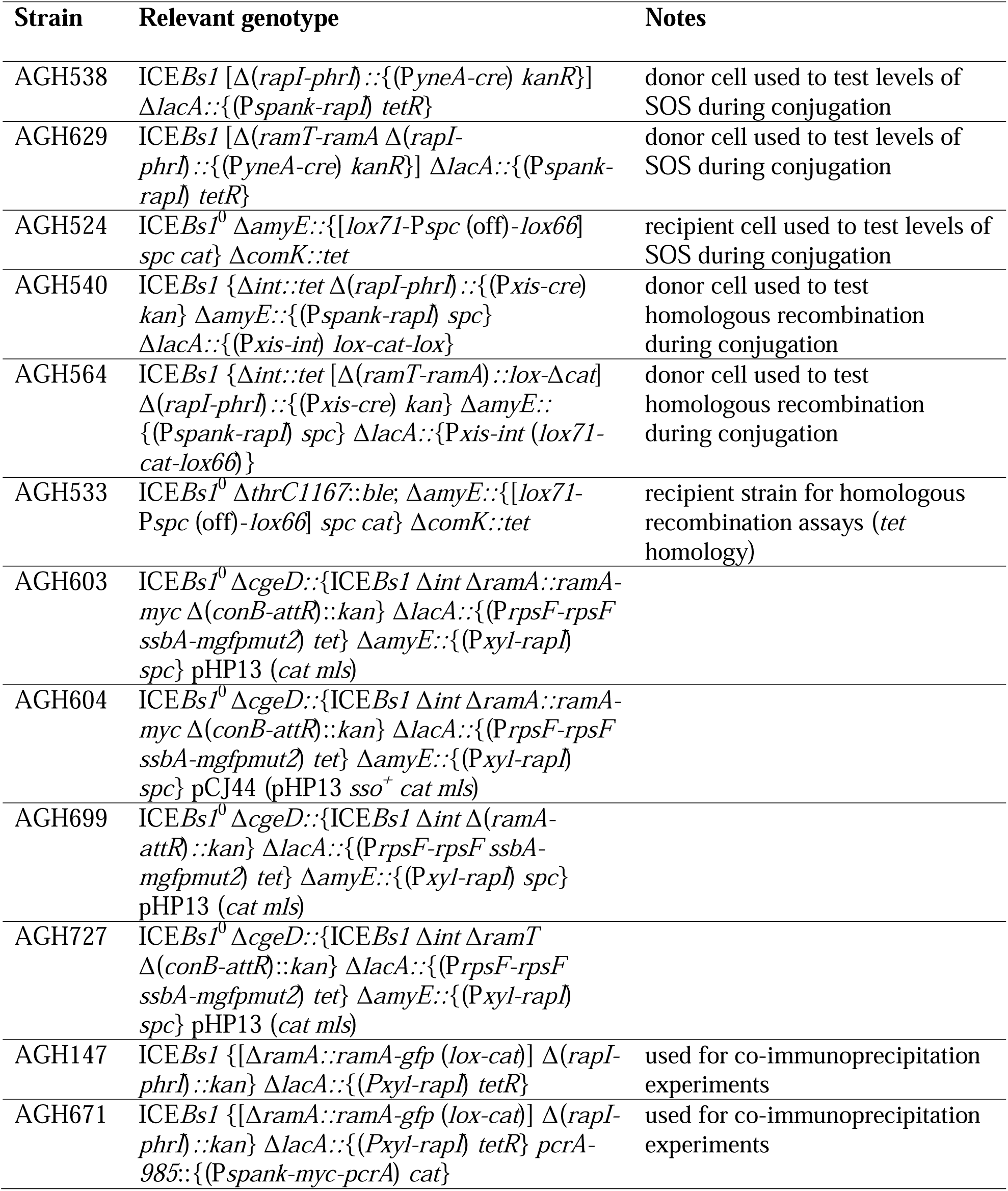
*B. subtilis* strains used.

#### PyneA-cre

P*yneA-cre* was used in conjugation experiments to test for SOS levels in recipient cells. Gibson isothermal assembly (Gibson et al., 2009) was used to piece together the promoter of the SOS-inducible *yneA* gene (155 bp that correspond the intergenic region between *laxA* and *yneA*) and *cre*, along with *kan* (kanamycin resistance) and flanking sequences for recombination into ICE*Bs1* at *rapI* and *phrI*. The homology flanking regions that delimited the *rapI* and *phrI* deletions where comprised from upstream homology flanking region contained the first 9 bp from *rapI* and extended 1000 additional bp upstream. This fragment was fused to another fragment containing the last 9 bp of *phrI* that extended an additional 1000 bp downstream. Transformation of this construct into competent *B. subtilis* generated strain AGH532. Genomic DNA from AGH532 was used to transfer Δ*rapIphrI::*P*yneA-cre* (*kan*) into subsequent strains to generate AGH540 and AGH564.

#### RamA-GFP

RamA-GFP was used in immunoprecipitation experiments. This fusion encodes a functional RamA with GFP at the C-terminus. The fusion gene was constructed using Gibson isothermal assembly of DNA fragments of *gfp*, *ramA*, *cat* (flanked by *loxP* sites), and flanking sequences that were generated by PCR. The assembled fragment was introduced into *ramA* by homologous recombination to generate SAM1089. Genomic DNA from this strain was used to transform *ramA-gfp* (*lox-cat-lox*) into subsequent strains to generate AGH147 and AGH671.

#### *ramT* and *ramA* deletions

Deletions of both *ramT ramA* (without an antibiotic resistance gene inserted) were made by using two different strategies. In the first, *cat* flanked by *loxP* sites was amplified and fused to sequences flanking *ramT* and *ramA* by Gibson isothermal assembly. This construct was originally defined in (McKeithen-Mead et al., 2024).

The second strategy to make deletions of *ramT* and *ramA* involved cloning the flanking regions into pCAL1422, a plasmid containing *lacZ* and *cat* in the backbone (Thomas et al., 2013), by Gibson isothermal assembly. More specifically, the homology flanking regions (which delimit the *ramT* and *ramA* deletions) comprised of an upstream homology flanking fragment that included the first 9 base pairs or *ramT* and extended 632 additional base pairs upstream *ramT*. This fragment was fused to another fragment that contained the last 39 bp from *ramA* and extended 1038 additional base pairs downstream *ramA*. The resulting plasmid (pAGH92) was used to transform appropriate *B. subtilis* strains, selecting for integration of the plasmid into the chromosome (chloramphenicol resistant) by single crossover recombination. Transformants were then screened for loss of *lacZ* and tested for deletions by PCR.

### Conjugation experiments

A single colony was inoculated into LB medium and grown at 37°C to mid-exponential phase. The culture was then diluted in pre-warmed LB to OD ∼ 0.01-0.025. Once the culture reached OD600 of 0.1, *rapI* was induced with 1 mM IPTG (from Pspank-*rapI*) for 40 min to induce ICE*Bs1*, after which mattings were set up in a 1:3 donor to recipient ratio. The cultures were vacuum-filtered, and the filters were incubated on Spizizen minimal salts plates (Auchtung et al., 2005) at 37°C for the indicated time. Then, the cells were collected from the filters in PBS and serial dilutions were performed and plated on agar plates containing appropriate antibiotics (kanamycin and chloramphenicol for the experiments that measured SOS levels; and kanamycin and phleomycin for experiments that tested for homologous recombination). These plates select for recipient cells that acquired ICE*Bs1* and became transconjugants. The conjugation efficiency was calculated as the number of transconjugant cells per the initial donor. One mL was saved for gDNA extraction for analysis by qPCR (see below). Each experiment was performed at least three times.

### Quantitative PCR analysis

We used pPCR to measure SOS levels in recipient cells during conjugation and to calculate the efficiency of homologous recombination. Both involved quantifying the frequency of inversion of P*spc* that was flanked by *loxP* sites. Expression of the Cre recombinase (either from the SOS promoter P*yneA*-*cre* or the ICE*Bs1* promoter P*xis*-*cre*) results the inversion of P*spc (McKeithen-Mead & Grossman, 2023)*. This Cre-mediated inversion was detected with the primers oSAM776 (5’-CCAGTCACGT TACGTTATTA GTTATAG -3’) and oSAM777 (5’-TACCGCACAG ATGCGTAAG -3’) (McKeithen-Mead & Grossman, 2023).

Recipients contained either *cat* or *ble* and these genes were used to measure the number of recipients by qPCR. Quantification of *cat* (amplified with primers LDW104 5’-GCGACGGAGA GTTAGGTTAT TGG-3’ and LDW107 5’-TTGAAGTCAT TCTTTACAGG AGTCC-3’) (Wright et al., 2015) and *ble* (amplified with primers AGH575 5’-ctacttcaatgcggcaactagc-3’ and AGH758 5’-gaagatggattcgcagttctaatgtg-3’) was used to determine the amount of SOS response in conjugation experiments and to calculate the ICE*Bs1* transfer efficiencies used to determine homologous recombination efficiencies, respectively.

### Yeast two hybrids

The *S. cerevisiae* strains used are derived from PJ69-4A (James et al., 1996) with previously described vectors (James et al., 1996). The entire coding sequence of *ramT* was cloned into pGAD and fused to the Gal4 activation domain, while the coding sequence of *ssbA* was cloned into pGBDU and fused to the Gal4 DNA binding domain. These vectors were transformed into competent *S. cerevisiae* cells using the LiAc protocol (Gietz & Schiestl, 2007). Transformants were plated on synthetic dropout medium with appropriate supplements to select for plasmid acquisition. Growth on agar plates lacking leucine or uracil indicated the acquisition of pGAD- and pGBDU-based plasmids, respectively. To test for protein interactions, yeasts carrying both plasmids were grown on plates lacking leucine, uracil, and adenine. Growth on these plates indicated an interaction between the fusion proteins. Yeast strains used included: CMJ620 (*S. cerevisiae* mat a ura3-52 leu2-3 his3 trp1 dph2Δ::HIS3 gal4Δ gal80Δ GAL2-ADE2 LYS2::GAL1-HIS3 met2::GAL-lacZ pCJ113 pCJ107); and AGH677 (*S. cerevisiae* mat a ura3-52 leu2-3 his3 trp1 dph2Δ::HIS3 gal4Δ gal80Δ GAL2-ADE2 LYS2::GAL1-HIS3 met2::GAL-lacZ pAGH646 pAGH651).

### ChIP experiments

Cells were grown on plates with appropriate antibiotics. A single colony was cultured overnight at 30°C in minimal media supplemented with L-arabinose as carbon source, and erthyromycin plus lincomycin to select for pHP13 plasmids. The following morning, cells were diluted to OD ∼0.025 in pre-warmed media supplemented with the antibiotics and grown to OD ∼0.2. RamT and/or RamA were induced by adding 1% xylose to activate *rapI* (Pxyl-*rapI*) for 90 minutes.

Chromatin immunoprecipitations were performed as previously described (Smits & Grossman, 2010; Merrikh & Grossman, 2011). Essentially, cells were crosslinked with formaldehyde (Sigma) at room temperature for 5 minutes, followed by quenching with glycine (0.25M final concentration) for an additional 5 minutes. Pellets were washed with PBS and lysed by sonication. Lysates were incubated overnight at 4°C with anti-GFP antibodies (A11120 ThermoFisher Scientific) and protein A beads (Cytiva Life Sciences). The next day, beads were washed, and protein-DNA complexes were eluted. SsbA-GFP binding to pHP13 was quantified by measuring the *cat* gene levels by qPCR, as pHP13 contains both *cat* and *mls* resistance cassettes (Wright et al., 2015). ChIP signals were normalized to total input for each sample. Each experiment was performed at least three times.

### Co-IP experiments

For co-immunoprecipitatin experiments, a single colony was inoculated into minimal medium supplemented with L-arabinose as carbon source and grown overnight at 30°C. The following morning, cells were diluted into pre-warmed medium to OD600 ∼0.02. At OD ∼0.2, ICE*Bs1* was induced by adding 1% xylose to activate *rapI* (Pxyl-*rapI*) for 90 minutes. Cells were harvested by centrifugation, washed with PBS, and lysed by sonication.

Immunoprecipitation was done essentially as described (Lin & Lai, 2017). Briefly, cell lysates were incubated with protein A beads (Cytiva Life Sciences) and anti-myc antibodies (132500, ThermoFisher Scientific) overnight at 4°C. Protein complexes bound to beads were washed and eluted and samples were resolved by SDS-PAGE on 12% polyacrylamide gels and transferred to nitrocellulose membranes (Cytiva Life Sciences). Primary antibodies used were anti-GFP (A11122 Invitrogen), and goat anti-rabbit (LICORbio) as the secondary antibody. Membranes were visualized using a Li-COR Odyssey imaging system (LICORbio).

## Acknowledgments

We thank members of the Grossman group, as well as Janeth Pérez-Garza for technical assistance and thoughtful discussions. Research reported here is based upon work supported, in part, by the National Institute of General Medical Sciences of the National Institutes of Health under award numbers GM122538 and GM148343 to ADG. Any opinions, findings, and conclusions or recommendations expressed in this report are those of the authors and do not necessarily reflect the views of the National Institutes of Health.

